# The nuclear localization of Ect2 is required for cytokinesis

**DOI:** 10.1101/2025.09.04.674117

**Authors:** Nhat Phi Pham, Gabrielle Schick, Joseph Del Corpo, Alisa Piekny

## Abstract

In cytokinesis, a contractile ring constricts the cell to form two daughters. After ingression, the ring matures into the midbody, and an intercellular bridge connects the daughter cells until it is cut during abscission. Ring assembly requires the activation of RhoA GTPase at the equatorial cortex by Ect2, a guanine nucleotide exchange factor (GEF). However, it is unclear if RhoA must be inactivated after ring closure to complete cytokinesis. Here, we show that the nuclear sequestration of Ect2 after ingression is required for cytokinesis. Mutating the nuclear localization signal (NLS) in Ect2 causes it to remain at the midbody where it generates persistent active RhoA, leading to cytokinesis failure. Re-localizing the mutant to the nucleus using SV40NLS restores cytokinesis. Further, over-expression of mutant Ect2 causes cytokinesis failure, which is rescued by reducing GEF activity. We propose that persistent RhoA causes instability in the intercellular bridge, preventing abscission. Nuclear sequestration may regulate the function of other contractile proteins with NLS sequences.

## Introduction

Cytokinesis is a highly complex process that occurs in multiple steps to physically separate a cell into two daughters at the end of the cell cycle (D’Avino et al., 2015; Basant and Glotzer, 2018; Verma and Maresca, 2019). The stages of cytokinesis must be carefully controlled to ensure that genetic material and cell fate determinants are equally allocated. Aberrant cytokinesis can result in the uneven distribution of these factors, causing chromosomal instability and contributing to developmental diseases and cancer (Lacroix and Maddox, 2012; Lens and Medema, 2019; Was et al., 2022). Cytokinesis begins with the assembly of a contractile ring by the activation of small GTPase RhoA by the GEF Ect2 (Piekny et al., 2005). Ect2 activation is spatiotemporally coordinated in anaphase by binding to RacGAP/Cyk4, part of the centralspindlin complex (Yüce et al., 2005; Kamijo et al., 2006; Wolfe et al., 2009). The ring then ingresses and transitions to a midbody ring, which surrounds the midbody at a bridge of microtubules that connects the two daughter cells (Echard et al., 2004; Öztop and Chaigne, 2025). The bridge is finally cut when the recruitment of the ESCRT-III machinery at the intercellular bridge triggers abscission, physically separating the daughter cells (Carlton et al., 2008; Mierzwa and Gerlich, 2014).

Given how rapidly a cell progresses through the earlier stages of cytokinesis, it is not fully understood how the ring undergoes dramatic changes to go from an actomyosin ring to the midbody, which is a distinct structure. Prior studies reported the shedding of membrane-bound septin-anillin-RhoA complexes as a mechanism to reduce their levels in the ring (El Amine et al., 2013). However, anillin also forms a separate complex with citron kinase that is retained and required to form a stable midbody (Kechad et al., 2012). Thus, anillin may be part of two independent complexes that modulate the cortex during the transition to forming a midbody ring and stable midbody (Kechad et al., 2012; El Amine et al., 2013; Carim et al., 2020). Actomyosin levels also decrease during ingression (Dambournet et al., 2011; Guizetti et al., 2011; Wang et al., 2019). Enzymes like cofilin depolymerize F-actin (Nagaoka et al., 1995), while the flavin monooxygenase MICAL is recruited to the newly formed midbody to further reduce F-actin via oxidation (Echard, 2008). Ect2 initially localizes to the midbody but then quickly re-localizes to the reforming daughter nuclei which could lead to a decrease in active RhoA (Tatsumoto et al., 1999).

It is not known if RhoA activity is required at the later stages of cytokinesis. Presumably some active RhoA is required for secondary ingression, which occurs before the bridge is finally severed (Schiel et al., 2011, 2012; Dema et al., 2018; Wang et al., 2019). In cells with lagging chromosomes, RhoA may also be required for cell elongation to promote clearing of chromatids from the bridge (Montembault et al., 2017). However, the majority of RhoA is likely removed or inactivated via mechanisms that include membrane shedding as mentioned above, via the decrease in Ect2 at the midbody, the delivery of p50RhoGAP-containing vesicles to the intercellular bridge, and other mechanisms that could include OSGIN, another flavin monooxygenase (Schiel et al., 2012; Goupil et al., 2024). The levels of active RhoA must be carefully controlled as excess active RhoA could lead to the retention of regulators that polymerize F-actin, leading to delayed abscission (Dambournet et al., 2011). A recent study showed that branched F-actin nucleated by Arp2/3 acts as a ‘plug’ to prevent the over-elongation of the ESCRT-III machinery in the intercellular bridge, which would prevent abscission by impairing the spastin-mediated severing of microtubules (Advedissian et al., 2024).

Ect2 may also need to be removed from the midbody for the recruitment of proteins controlling midbody maturation and abscission. A previous study showed that as Ect2 leaves the midbody, Cyk4 transitions to forming new complexes with Fip3, which could recruit recycling endosomes required for abscission (Simon et al., 2008). After ingression, the GTPase Rab11 and its effector Fip3 deliver vesicles to the intercellular bridge, and this is required to stabilize and narrow the intercellular bridge (Mierzwa and Gerlich, 2014). Ect2 may compete with Fip3 for Cyk4-binding, as overexpression of an Ect2 fragment that binds to Cyk4 inhibits Fip3 recruitment to the midbody (Simon et al., 2008). However, the timing of Ect2’s localization at the midbody is not clear as there have been no spatiotemporal studies with high resolution done in human cells to properly characterize endogenous Ect2 during the later stages of cytokinesis.

Whether Ect2 needs to be reduced at the midbody for a decrease in active RhoA or for new Cyk4-complexes, nuclear import could act as a removal mechanism. Nuclear import already plays a significant role in the spatiotemporal regulation of mitosis. Prior to mitosis, spindle assembly factors are localized in the nucleus, then released during prometaphase to form the mitotic spindle (Kalab and Heald, 2008; Heald and Khodjakov, 2015). In interphase, these factors are sequestered in the nucleus by importins, which bind to their nuclear localization signals (NLS) (Cavazza and Vernos, 2016; Lu et al., 2021). Since several essential cytokinesis regulators have NLSs and localize to the nucleus in interphase, nuclear sequestration is likely to play a regulatory role. Deletions that block the nuclear localization of Ect2 cause multinucleation, while the overexpression of a cytosolic N-terminal fragment causes a gain-of-function phenotype and induces malignant transformation in cultured cells (Tatsumoto et al., 1999; Saito et al., 2004; Chalamalasetty et al., 2006). However, these studies were either done before there were extensive tools available for live-cell imaging studies, and/or in the presence of endogenous Ect2, and it is not clear how mutating the NLS disrupts Ect2 function. Mis-localized Ect2 is also associated with poor prognosis in patients with colorectal or lung cancers, although the mechanism is unknown. Similarly, a NLS mutant of Mklp1/Kif23, which forms the centralspindlin complex with Cyk4, is unable to rescue the depletion of endogenous protein, leading to cell multinucleation (Liu and Erikson, 2007). Although mutating the N-terminal NLS of anillin does not lead to cytokinesis failure in HeLa cells, cytosolic anillin causes abnormal cell shape changes (Chen et al., 2015). We hypothesize that the nuclear re-sequestration of contractile proteins after ring ingression may play a significant role in regulating the later stages of cytokinesis as a mechanism to control midbody maturation and/or abscission, or for G1-specific processes.

Ect2 has a functional NLS that controls its nuclear import in interphase cells. Whether the pool of Ect2 in the newly formed daughter nuclei is the result of re-localized Ect2 from the midbody or from *de novo* translation is unknown. Ect2 has TEK and D-box domains that target the protein for degradation by the APC^Cdh1^ (Liot et al., 2011). Abolishing nuclear localization impairs Ect2 degradation, suggesting that Ect2 re-localizes from the midbody to the nucleus after ingression where it is targeted for degradation. Nuclear import could therefore act as a mechanism to remove Ect2 from the midbody and degrade it in the nucleus. Since there are no temporal studies of Ect2, it is not known how long Ect2 remains at the midbody for, and/or if any pools remain for the later stages of cytokinesis.

Here, we show that the removal of Ect2 from the midbody by nuclear import is required for cytokinesis. Mutating the NLS causes Ect2 to remain at the midbody for much longer than wild-type Ect2 and leads to failed cytokinesis in Ect2-depleted cells. In these cells, the intercellular bridge is unstable and blebs, and ring closure is partially impaired as seen with endogenous anillin localization. Cytokinesis failure caused by the NLS mutant can be rescued by adding a SV40NLS sequence at the C-terminus. Using the active RhoA reporter 2xrGBD, we show that active RhoA persists for longer in cells expressing the Ect2 NLS mutant. Overexpressing Ect2 with the NLS mutant causes an increase in cytokinesis failure, and mutating the DH-domain (essential for RhoA activity) rescues this phenotype, suggesting that the failure was caused by excess RhoA activity. Together, these results support a model where nuclear import actively removes Ect2 after ingression to reduce RhoA activity and form a stable intercellular bridge.

## Materials & Methods

### Cell Culture and Transfection

HeLa cells were plated and grown in DMEM (Wisent), supplemented with 10% cosmic calf serum (Thermo Scientific) and were maintained at 37°C with 5% CO_2_. 5×10^5^ cells plated in 2mL media prior to transfection. Cells were transfected using Lipofectamine 3000 according to the manufacturer’s protocol (Invitrogen). Briefly, 2.5 μL of Lipofectamine was used per 2 mL of media with 2.5 μg DNA, 2.5 µL of P3000 and 2.5 nM siRNAs, as described previously (Yüce et al., 2005; Piekny and Glotzer, 2008). Cells transfected with plasmids encoding both mScarlet-tagged proteins and shRNAs were imaged 24–48 h after transfection, whereas cells co-transfected with plasmid DNA and siRNAs were fixed after 30 h. HeLa cells endogenously-tagged with mNeonGreen at the anillin, Ect2 or RhoA loci were previously generated (Husser et al., 2022).

### Constructs

All plasmids were maintained and cloned in Escherichia coli DH5α unless specified otherwise. The myc:Ect2 constructs were previously generated (Yüce et al., 2005). mScarlet-I-Ect2 was generated by first cloning mScarlet-I, amplified from mScarlet-I-mTurquoise2 (Addgene #98839; Mastop et al., 2017), and Ect2, amplified from myc:Ect2 (Yüce et al., 2005) into the pYTK001 backbone (Addgene #65108; Lee et al., 2015) by Golden Gate using a standard protocol, then assembled into a modified pX459V2.0-HypaCas9 vector (Addgene #108294; Kato-Inui et al., 2018) for mammalian expression. The shRNA sequence targeting endogenous Ect2 was the same as the siRNA sequence described in Yüce et al., (2005) and was cloned into the modified pX459V2.0-HypaCas9 vector beforehand. mNG-2xrGBD was generated by amplifying 2xrGBD from dTomato-2xrGBD (Addgene #129625; Mahlandt et al., 2021) and mNeonGreen (Allele Biotech) by Golden Gate cloning into pYTK001 followed by a modified pX459V2.0-HypaCas9 vector as previously described. Restriction enzymes used for Golden Gate assembly were BbsI, BsmBI-v2 and BsaI-hfv2 (New England Biolabs). Substitutions (Ect2 347AAAAA351, Ect2 348AAA350, Ect2 565AAA568 + P570S), insertions (Ect2 348AAA350 + SV40NLS) and truncations (Ect2 328-388) were generated using a one-step PCR protocol with partially overlapping primers (Liu and Naismith, 2008). GST:importin-β was previously generated (Beaudet et al., 2020). All constructs were verified by sequencing (Genome Quebec, Plasmidsaurus, and FlowGenomics).

### Cell fixation and Immunostaining

To fix cells for imaging, they were seeded to 50% confluency on 22×22mm glass coverslips previously etched with 100 mM HCl. 30 hours after co-transfection of DNA and siRNAs, the cells were fixed in fresh, ice-cold 10% w/v trichloroacetic acid for 14 minutes before being washed three times with PBST (1 X PBS with 0.5% Triton X-100) as previously described (Yüce et al., 2005)\.

For immunostaining, cells were incubated in normal donkey serum (NDS; 5% in PBST) for 20 minutes. After blocking, primary and secondary antibodies were added to the fixed cells for 2 hours each, washing with PBST in between. Myc was stained with primary anti-Myc monoclonal antibodies (1:250; DSHB) and anti-mouse Alexa-568 (Invitrogen; 1:500), and anillin with primary anti-anillin rabbit polyclonal antibodies (1:250; Piekny and Glotzer, 2008) and secondary anti-rabbit Alexa-488 antibodies (Invitrogen; 1:400). After the antibodies, cells were incubated with DAPI (1/1000 dilution from 1mg/mL stock; Sigma) for 5 minutes. Coverslips were mounted onto glass slides with mounting media (4% n-propyl gallate in 50% glycerol diluted in 50 mM Tris pH 9 and water) and sealed with nail polish to prevent drying.

### Microscopy

Fixed cells transfected with Myc::Ect2 and co-stained for anillin and DAPI were imaged using a Leica DMI6000B wide-field microscope with the 63×/1.4 PL APO oil immersion objective (pixel size 0.102 μm), and Z-stacks of 0.3 μm were acquired with a Hamamatsu OrcaR2 camera and Volocity software (PerkinElmer) using a piezo Z stage (MadCityLabs). Image files were exported as TIFFs, which were opened with Fiji (National Institutes of Health; NIH) and converted into maximum intensity Z-stack projections. Projections and merged color images were then converted into 8-bit images and imported into Illustrator (Adobe) to make figures. To perform live imaging, cells were plated and transfected on 25-mm round coverslips (no. 1.5) or 22×22-mm square coverslips (no 1.5) and incubated with Hoechst 34580 (Invitrogen) at 0.89 μM or SYTO Deep Red (Invitrogen) at 50 nM for 30 min and placed in a Chamlide magnetic chamber (Quorum). Cells were kept at 37°C with 5% CO_2_. Live imaging was performed on an inverted Nikon Eclipse Ti microscope with a Livescan Swept Field confocal unit (Nikon), using the 100×/1.45 CFI PLAN APO VC oil immersion objective (Nikon), a piezo Z stage (MadCity Labs), and with the Evolve EMCCD camera (Photometrics). Images were acquired using the 405-nm, 561-nm and 640-nm lasers (100 mW; Agilent) set between 3 and 15% power, depending on the intensity of fluorescent signals (settings were kept constant for related experiments), and multiple Z-stacks of 1.0 μm were taken every 120 s per cell using NIS-Elements acquisition software (Nikon), and a quad filter (430-485 + 520-550 + 590-630 +680-740; Chroma) or far-red filter. Cells were also imaged on an inverted Zeiss Axio Observer with a Cicero spinning disk (CrestOptics), using a 63×/1.4 PLAN APO DIC M27 oil immersion objective (Zeiss; pixel size 0.26 μm), with a piezo Z stage (MadCityLabs), and an Orca Flash4.0 LT camera (Hamamatsu). Images were acquired using the 555-nm and 637-nm lasers (89North) set between 15-20% power. Multiple Z-stacks of 1.0 μm were taken every 120 s per cell for 60 min then every 10 min per cell for 3 hours, using Volocity (Quorum). Cells were also imaged on an inverted Nikon Eclipse Ti2 microscope with a CSU-X1 spinning disk (Yokogawa), using the 60×/1.4 CFI PLAN APO VC oil immersion objective (Nikon), with a piezo Z stage (MadCityLabs), and with the Zyla camera (Andor). Images acquired using the 561-nm and 638-nm lasers (OMICs) set at 3% power, and multiple Z-stacks of 1.0 μm were taken every 120 s per cell, or 120 s per cell for 60 min then every 10 min for 3 hours using μManager (NIH). Image files were exported as TIFFs, which were opened with Fiji (NIH) and converted into maximum intensity Z-stack projections. Projections and merged color images were then converted into 8-bit images and imported into Illustrator (Adobe) to make figures.

### Pull-down assays

The purified proteins GST and GST:importin-β1 were made from Escherichia coli BL21 cells. Bacteria were resuspended in lysis buffer (2.5 mM MgCl_2_, 50 mM Tris, 150 mM NaCl, pH 7.5, 0.5% Triton X-100, 1 mM dithiothreitol [DTT], 1 mM phenylmethanesulfonyl fluoride [PMSF], and 1× protease inhibitors [Roche]), incubated with 1 mg/mL lysozyme on ice for 30 min, then sonicated three times. Extracts were incubated with preequilibrated glutathione sepharose 4B (GE Lifesciences) overnight at 4°C with rotation. After washing, beads were stored as a 50% slurry at 4°C. Protein concentration was determined by running samples by SDS–PAGE and measuring the density of bands in comparison to known concentrations of bovine serum albumin. To test for binding, proteins were pulled down from cell lysates after transfection. Transfected HeLa cells were lysed in 50 mM Tris, pH 7.6, 150 mM NaCl, 5 mM MgCl_2_, 0.5% Triton X-100, 1 mM DTT, 1 mM PMSF with 1× protease inhibitors (Roche), and incubated with 5–10 μg of purified GST-tagged importin-β protein on beads at 4°C overnight. After binding, beads were washed three to four times with 50 mM Tris, pH 7.6, 150 mM NaCl, 5 mM MgCl_2_ before adding SDS sample buffer to denature the proteins for SDS PAGE. All samples were run by SDS–PAGE and wet-transferred to nitrocellulose membrane for Western blotting. All blots were reversibly stained with Ponceau S to show total protein. The blots were blocked with 5% w/v milk for 20 min, then incubated with 1:10,000 mouse anti-mNeonGreen antibodies (ChromoTek) in 1× TBS-T (150 mM NaCl, 100 mM Tris pH 7.4, 0.5% Triton X-100) for 1–2 h at room temperature. After washing the membrane three to four times with 1× TBS-T, secondary antibodies (anti-rabbit HRP [horseradish peroxidase]; New England Biolabs) were added as per manufacturer’s instructions in 1 × TBS-T for 2 h. The blots were developed using enhanced chemiluminescence (ECL) Western blotting detection reagents (Cytiva) and visualized on a GE Amersham Imager 600 or ChemiDoc XRS+ (Bio-Rad). All results from each pull-down assay were replicated at least three times to ensure reproducibility. Images were converted to 8 bit by Fiji, and made into figures using Illustrator (Adobe).

### Analysis

All images acquired using NIS Elements (Nikon) were opened in Fiji (Version 2.3, NIH) or IMARIS (Oxford Instruments) for analysis. Linescans were performed and measured using a macro in Fiji modified from Ozugergin et al., (2022). The macro was designed to isolate the desired channel from the image file, subtract background signal, and perform a bleach correction. The desired timepoint and three central Z slices were picked manually, and the macro generated a Z-stack sum-slice projection. A five-pixel-wide line was then traced along the cortex of the cell, from one pole to the other, along with a straight one-pixel-wide line to define the midplane. The macro then measured the fluorescence intensity of each pixel along the length of the linescan and positioned the pixels in relation to the midplane. For breadth measurements, the number of pixels above 50% of the normalized peak intensities were counted for each linescan and converted to microns. Pixels with intensities higher than the cutoff value outside of the peak region were excluded from these calculations. To measure the ratio of cortical accumulation versus pole, the area under the curve of the fluorescence intensity delineated by the breadth was divided by the area under the curve of the same for the average of the first and last 10% pixels of the linescan. Kymographs were performed and measured using a macro in Fiji. The macro was designed to isolate the desired channel, and a five-pixel-wide line was drawn manually over the furrow region at every timepoint until closure. Then, the distance between the two sides of the membrane was measured at each timepoint using the straight-line tool, and measurements were exported to Excel (Microsoft). The distance between the two sides at anaphase onset was set to a maximum value (100%) and used to normalize the distance throughout ingression. As the closure times were variable among cells, measurements were terminated when at least three cells had completed cytokinesis. For measurement of fluorescence intensity over time, images were analyzed using Imaris. The signal at the intercellular bridge or midbody was modeled as a spot or surface and tracked over time and its sum of fluorescence intensity over time was determined. For the ratio of sum-intensity at closure and before closure, the sum intensity of fluorescence intensity at closure was divided by the same at the timepoint before closure. All data were imported into Excel (Microsoft) for formatting, then to Prism (Version 9.3, GraphPad) for further analysis. All images and graphs were transferred to Illustrator (Adobe) to make figures.

## Statistical analysis

Line graphs, bar graphs and box-and-whisker plots were generated using Prism (Version 9.3, GraphPad). Distribution of the data was assessed, and statistical significance was tested using Student’s t-test or a one-way ANOVA followed by multiple comparisons using a Tukey’s post-hoc test if the data was normally distributed, or using a non-parametric test if the data was not normally distributed. Significance levels were defined as: p>0.05 non-significant (ns), *p≤0.05; **p≤0.01.

## Results

### Endogenous Ect2 localizes to the midbody and daughter cell nuclei after ring ingression

Ect2 localization was previously reported to change throughout the cell cycle (Tatsumoto et al., 1999). In interphase cells, Ect2 is in the nucleus, then becomes cytosolic after nuclear envelope breakdown where it localizes to the central spindle and midbody (Yüce et al., 2005; Simon et al., 2008). However, since these studies relied on fixed cells, the timing of changes in Ect2 at the midbody was not known, or if this reduction occurred because of its transport to the daughter cell nuclei or from degradation (Liot et al., 2011). Using HeLa cells where Ect2 is endogenously tagged with mNeonGreen, we imaged Ect2 with high spatiotemporal resolution throughout cytokinesis, including abscission (Figure 1B; Husser et al., 2022). As expected, Ect2 localized to the central spindle and overlying cortex, followed by localization to the midbody and daughter cell nuclei. The increased temporal resolution of our data revealed that Ect2 started to appear in the daughter cell nuclei shortly after ring closure, and there was a decrease at the midbody with concomitant increase in the daughter cell nuclei (Figure 1B). We quantified the change in Ect2 intensity at the midbody and nuclei over time starting at ring closure (t = 0; Figure 1C). As shown in the graph, there was a steady decrease in Ect2 at the midbody, with the exception of a slight increase at t = 20 minutes, which was no longer visible by 100 minutes, and corresponded with an increase in nuclear Ect2 (Figure 1C). This suggests that the nuclear transport of Ect2 from the midbody reduces its levels there, and the nuclear pool of Ect2 is likely degraded >100 minutes after ring closure.

**Figure 1.**
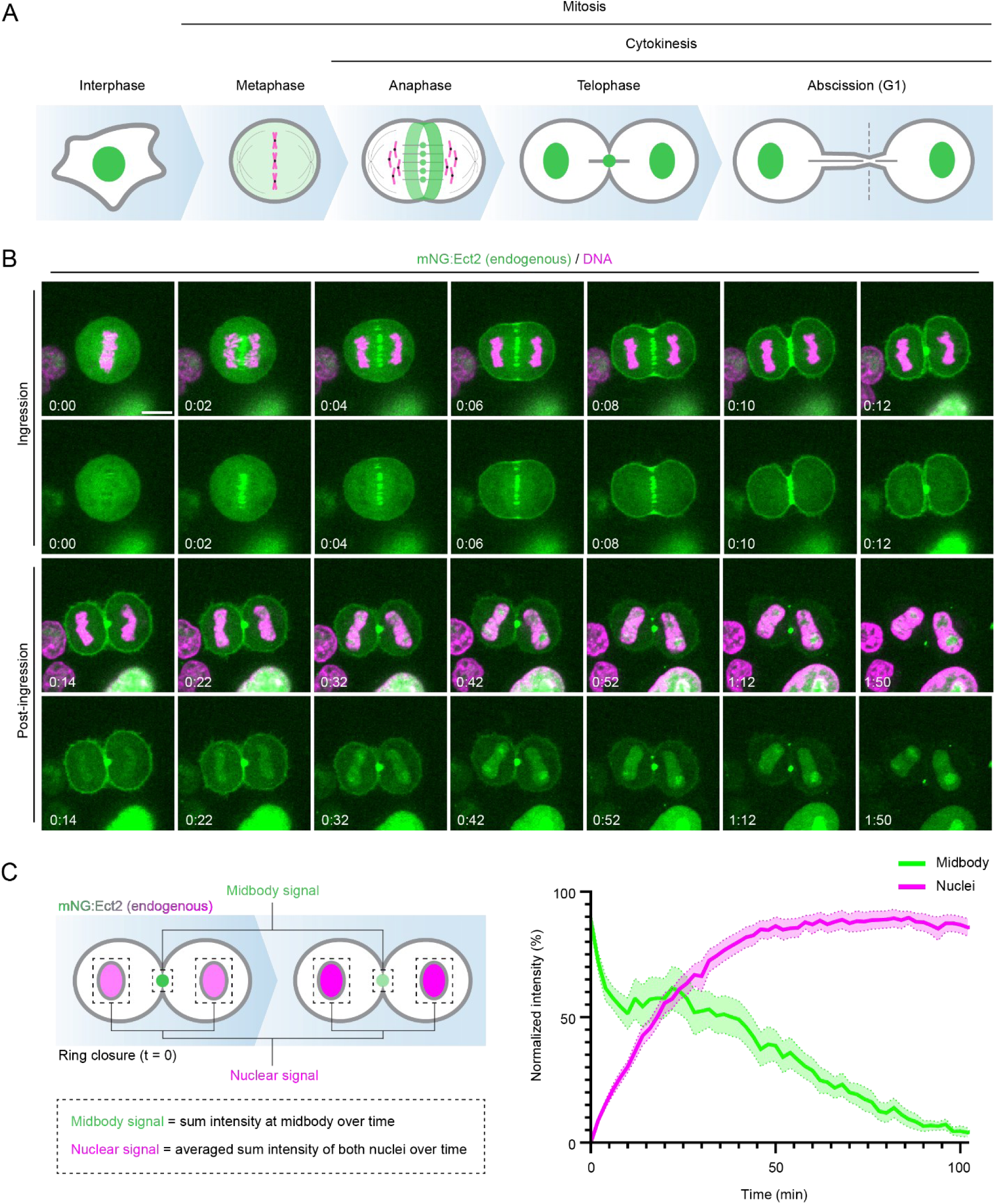
Ect2 decreases at the midbody with concomitant increase in the daughter cell nuclei. (A) Cartoon cells show Ect2 localization (green; chromosomes are in pink) in the nucleus in interphase, and throughout mitosis where it transitions from being cytosolic, to the central spindle and ring, then to the midbody and nuclei of the daughter cells. (B) Timelapse images show a HeLa cell expressing endogenous mNeonGreen (mNG):Ect2 (green) co-stained for DNA (magenta, SYTO Deep Red). Images taken after ring closure are labeled as “post-ingression”. The scale bar is 10 μm. Time is in hours:minutes (t = 0 minutes refers to anaphase onset, and ring closure is at 12 minutes). (C) The schematic on the left shows how the midbody (green) and nuclear pools (magenta) of Ect2 were quantified by measuring the sum total fluorescence intensity over time after ring closure (note that the time t = 0 now refers to ring closure). On the right, a line graph shows a plot of the normalized fluorescence intensity (y-axis, %) of the midbody (green) and nuclear pool (magenta) over time (x-axis, minutes; n = 14 cells). The error bars show standard error of the mean (SEM).

### The nuclear localization of Ect2 is required for cytokinesis

Previous studies showed that the S-loop NLS is required for nuclear localization, however, it was not known if Ect2’s nuclear localization is required for cytokinesis. To test this, we mutated the NLS of Ect2 by substituting 348KRR350 to alanine residues (348KRR-AAA350; Ect2^3A^) to prevent nuclear transport (Figure 2A). The nuclear localization of mScarlet-I-Ect2^3A^ was disrupted in interphase HeLa cells compared to the non-mutant control (Figure 2B). We also observed a reduction in the binding of mNeonGreen:Ect2^3A^ (328-388) from HeLa cell lysates to recombinant Glutathione S-transferase (GST)-tagged importin-β1 compared to the non-mutant control (Figures 2C, D). Next, we determined if the NLS is required for cytokinesis by performing rescue assays by expressing Myc-tagged RNAi-resistant Ect2^3A^ or a non-mutant control in cells co-depleted of endogenous Ect2 using siRNAs (Figure 2E). Since cytokinesis failure causes cells to become binucleate, we used the number of binucleate cells as a measure of cytokinesis failure (Figure 2E). While 63.5% of cells were binucleate after Ect2 RNAi (control), 17.7% of cells rescued with RNAi-resistant non-mutant Ect2 were binucleate, and 49.2% of cells rescued with RNAi-resistant Ect2^3A^ were binucleate (Figure 2F). Therefore, the NLS of Ect2 is required for cytokinesis. We also observed that the over-expression of Ect2 without depletion of endogenous Ect2 caused 10.5% of cells to become binucleate, which is consistent with previously published data (Figure 2F; Hara et al., 2006). Interestingly, over-expressing Ect2^3A^ caused a significant increase in the proportion of binucleate cells (37.1%) compared to non-mutant Ect2 suggesting that the NLS mutant is also dominant-negative. We observed a similar effect in HEK293T cells, which are non-cancerous. While the over-expression of Ect2 did not cause cytokinesis failure on its own, over-expression of Ect2^3A^ caused 76.9% of cells to be binucleate (Figures S1A, B). This data suggests that the requirement for nuclear localization is conserved. Furthermore, the levels of Ect2 in the cell are critical for its function in cytokinesis, and nuclear localization could be a mechanism to manage these levels.

**Figure 2.**
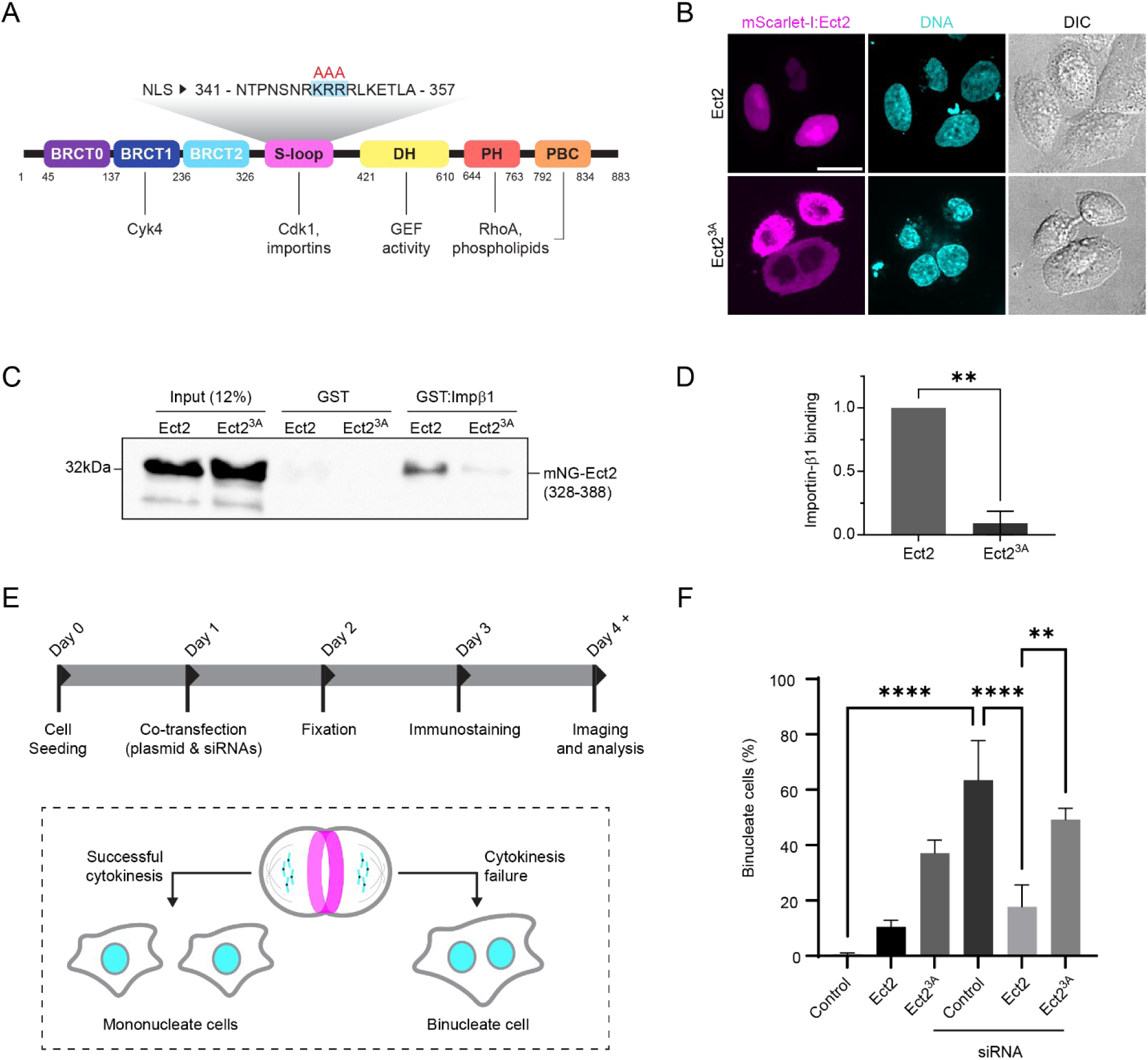
The NLS of Ect2 is required for cytokinesis. (A) A cartoon structure of Ect2 is shown with binding domains as indicated. The BRCT domains bind to Cyk4/MgcRacGAP. The S-loop contains a Cdk1 phosphorylation site, and an NLS (sequence shown above with the mutations using in this study in red) that binds to importins and controls nuclear localization. The Dbl-Homology (DH) has GEF activity for RhoA, while the Pleckstrin Homology (PH) and PolyBasic Cluster (PBC) regions bind to phospholipids. (B) Images show HeLa cells expressing mScarlet-I:Ect2 (magenta; Ect2 or Ect2^3A^), co-stained for DNA (cyan, Hoechst) and imaged with DIC (grayscale). The scale bar is 10 μm. (C) A western blot shows the binding of mNeonGreen (mNG)-tagged Ect2 (328-388; Ect2 or Ect2^3A^) from HeLa cell lysates to purified recombinant GST:importin-β1 (GST:Impβ1). (D) A bar graph shows the mean densitometry measurements, normalized to input and to the control (Ect2, WT). Error bars show standard deviation (N = 3). Statistical analysis was done using Student’s T-test (**, p<0.01). (E) A timeline shows the experimental design for the rescue assay and a cartoon shows the consequences of successful cytokinesis or failure. (F) A bar graph shows the proportion of binucleate cells as a measure of failed cytokinesis (y-axis, %) in HeLa cells expressing RNAi-resistant myc:Ect2 (Ect2 or Ect2^3A^) +/-Ect2 siRNA. Error bars show standard deviation (N = 3, with n = 26-249 cells per replicate). Statistical analyses were done using ANOVA and Tukey post-hoc test (ns, not significant; ***, p<0.0002; ****, p<0.0001).

### The nuclear localization of Ect2 controls its levels at the midbody

To determine the cytokinesis phenotypes caused by the Ect2^3A^ mutant, we performed live-cell imaging. HeLa cells expressing RNAi-resistant mScarlet-I-tagged Ect2 or Ect2^3A^ co-depleted of endogenous Ect2 were imaged throughout cytokinesis (Figure 3A). During mitotic exit, Ect2 localized to the spindle midzone and equatorial cortex, then to the midbody and subsequently to the daughter nuclei as shown in Figure 1, and the majority of cells completed cytokinesis successfully (10%; Figures 3A, B). Ect2^3A^ also localized to the spindle midzone and equatorial cortex, and subsequently to the midbody. However, the mutant remained at the midbody, and in 44.2% of the cells, the membrane regressed causing the cells to become binucleate (Figures 3A, B). We quantified the levels of Ect2 at the midbody from ring closure until it was no longer detectable or before regression (Figure 3C). While Ect2 decreased until it was no longer detected at 70 min after closure, Ect2^3A^ remained at the midbody for a longer period of time and persisted until membrane regression (Figure 3C). These results suggest that cytokinesis failure could arise because Ect2 remains at the midbody after ring closure, rather than being transported to the daughter cell nuclei. To test this, we fused the SV40NLS to the C-terminus of Ect2^3A^ (Ect2^3A-SV40NLS^) and expressed the mScarlet-tagged RNAi resistant mutant in cells co-depleted for endogenous Ect2. As shown in Figure 3A-C, adding the SV40NLS restored the nuclear localization of Ect2^3A^, which decreased at the midbody like non mutant Ect2 and rescued cytokinesis (16.2% binucleate cells). This data, summarized in Figure 3D, supports that the nuclear localization of Ect2 from the midbody is required for cytokinesis. Interestingly, Ect2^3A-SV40NLS^ localizes to the daughter cell nuclei earlier than Ect2, indicating that the rate of nuclear transport could vary with the NLS sequence or their position in the protein.

**Figure 3.**
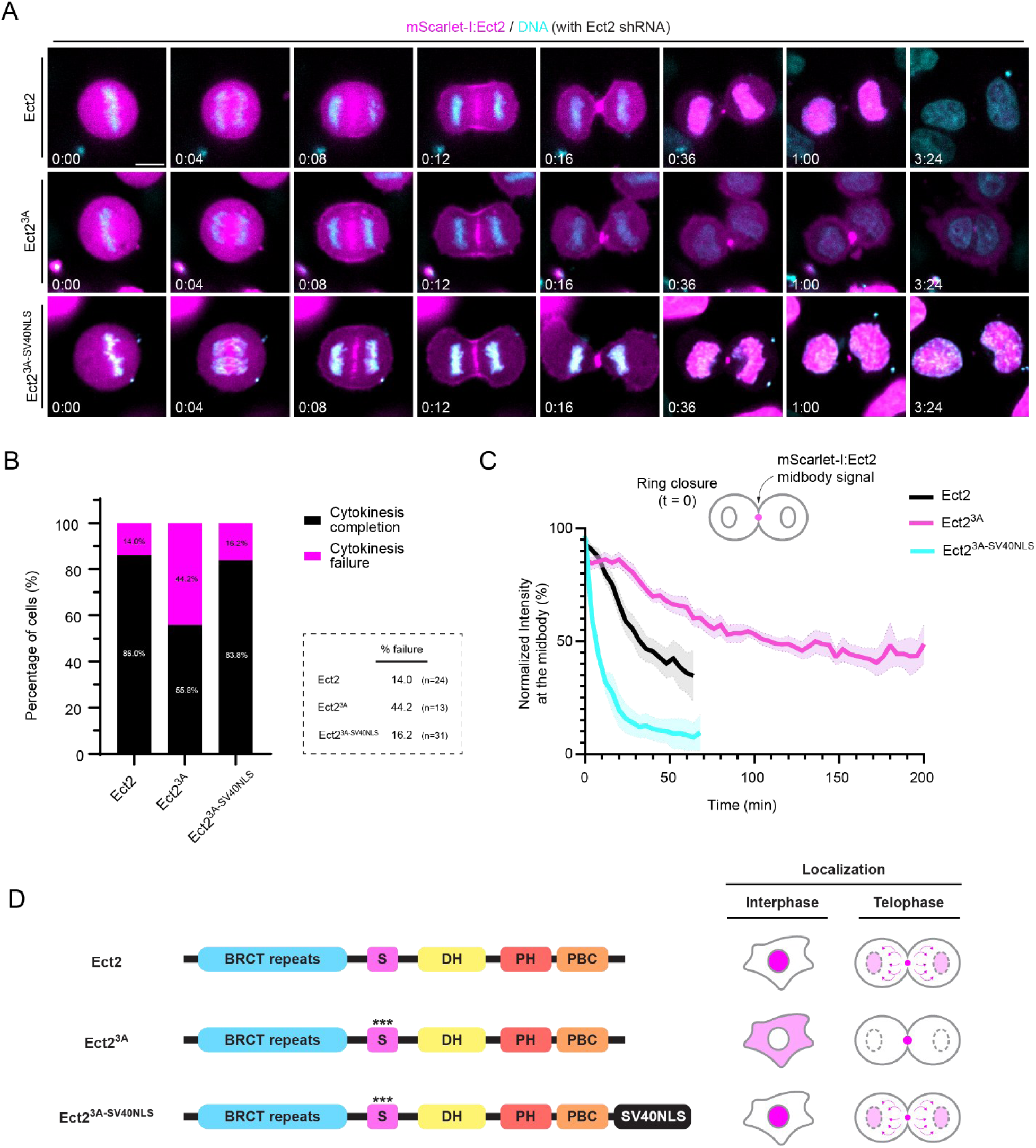
The nuclear localization of Ect2 is required to reduce its levels at the midbody. (A) Timelapse images show HeLa cells expressing RNAi-resistant mScarlet-I-Ect2 (magenta; Ect2, Ect2^3A^, Ect2^3A-SV40NLS^) depleted of endogenous Ect2 using shRNA, co-stained for DNA (cyan, SYTO Deep Red). The scale bar is 10 μm. Time is in hours:minutes (t = 0 is anaphase onset). (B) A bar graph indicates the proportion of cells that succeed (black) and fail (pink) cytokinesis for cells as shown in (A; Ect2 14% failure, n = 24; Ect2^3A^ 44.2% failure, n = 13; Ect2^3A-SV40NLS^ 16.2% failure, n = 31). (C) A line graph shows the changes in normalized fluorescence intensity (y-axis, %) at the midbody over time (x-axis, minutes in cells as shown in A (Ect2, black, n = 10; Ect2^3A^, pink, n = 15; Ect2^3A-SV40NLS^, blue, n = 11). The cartoon above shows how the midbody intensity was quantified. The error bars indicate standard error of the mean (SEM). (D) Ect2 structures with the NLS mutations (***) and SV40NLS (black) in A-C are shown (left) along with their localization in cartoon cells in interphase and telophase (right).

We also determined if Ect2^3A^ causes earlier defects in the contractile ring since earlier problems with ring assembly or ingression could impact midbody formation. We noticed that in some cells that were rescued with Ect2^3A^ the ring appeared to take longer to close compared to Ect2. We measured ingression rates using kymographs of the equatorial plane to quantify the extent of ring closure, then plotted the changes in ring diameter over time (Figure S2A). Indeed, there was a slight, but measurable delay in the closure of the ring in cells rescued with Ect2^3A^ compared to Ect2 (Figure S2B). While in cells rescued with Ect2, the ring closed ∼10-12 min after anaphase onset, it took ∼14-16 min in Ect2^3A^ cells. To determine if this delay was due to defects in ring assembly, we measured changes in the recruitment and breadth of Ect2 or the ring protein anillin, where anillin was endogenously tagged with mNeonGreen. To do this, the fluorescence intensity was measured along the cortex, then the breadth was measured as the number of the pixels > 50% of maximum fluorescence intensity, which was normalized by calculating the ratio vs. the length (total number of pixels; Figure 3A). The intensities of Ect2^3A^ were more variable compared to Ect2, but the breadth of Ect2 accumulation in the equatorial plane was similar (Figure S3B), while the intensities and breadth of anillin were similar in cells rescued with Ect2 or Ect23A (Figure S3C). Since anillin recruitment depends on active RhoA, Ect2^3A^ can generate sufficient active RhoA for ring assembly.

### Midbody formation requires Ect2 removal

Our data supports that Ect2’s nuclear localization is required for successful cytokinesis, and its persistence at the midbody causes cytokinesis failure. To determine why, we characterized the localization of anillin, which was previously shown to be required for the ring to midbody transition (Kechad et al., 2012). While endogenous mNG anillin localization was similar in cells rescued with Ect2^3A^ compared to Ect2 during ring assembly and ingression, we observed a difference after ring closure (Figure 4A). Anillin was more broadly localized and/or intense, and blebs appeared around the intercellular bridge (Figure 4A). To quantify this phenotype, we measured changes in the ratio of anillin fluorescence intensity at closure (when a change in ring diameter cannot be detected) compared to the timepoint just before closure, which could indicate perturbations in the ring-to-midbody transition (Figure 4B). Cells rescued with Ect2^3A^ showed a greater range and variability in anillin compared to Ect2, suggesting that midbody formation is impaired in the mutant (Figure 4B).

**Figure 4.**
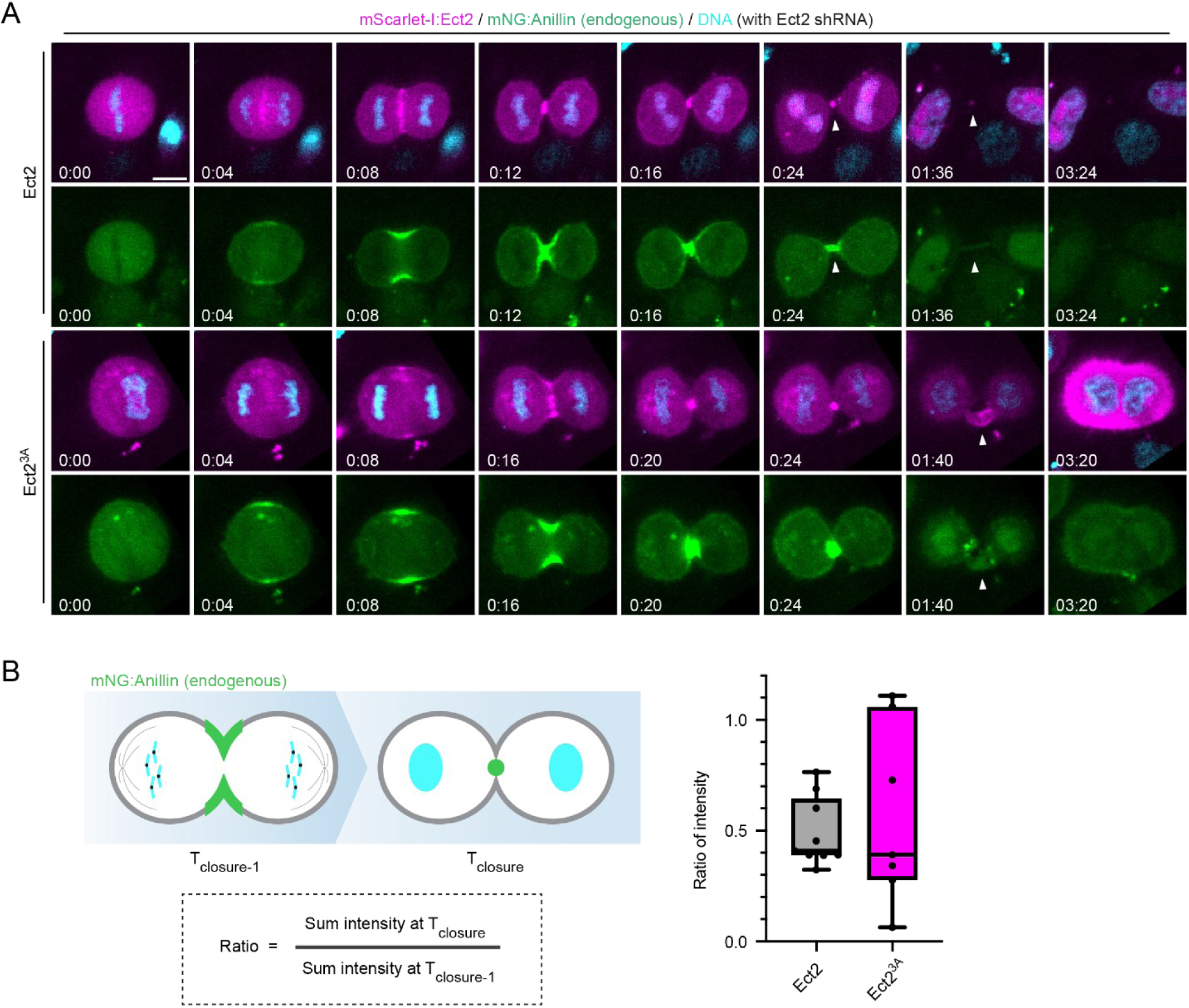
Removal of Ect2 from the midbody is required for its formation. (A) Timelapse images show endogenous mNeonGreen:anillin (green) in HeLa cells co-expressing RNAi-resistant mScarlet-I-Ect2 (magenta; Ect2 or Ect2^3A^), depleted of endogenous Ect2 using shRNA and co-stained for DNA (cyan, Hoechst). The scale bar is 10 μm (t = 0 is anaphase onset). Time is in hours:minutes. White arrowheads point to the intercellular bridge. (B) On the left, cartoon cells indicate how the ratio of sum anillin intensity at the midbody was measured at ring closure (T_closure_) vs. just before closure (T_closure-1_). On the right, a box-and-whiskers plot shows the ratio of anillin intensities (y-axis) in cells rescued with mScarlet-I:Ect2 (grey, n = 9) and Ect2^3A^ (pink, n = 7) as imaged in (A). Bars indicate standard deviation, and the black line shows the mean.

### Ect2 removal causes a decrease in active RhoA for midbody formation

Our data supports that Ect2 nuclear localization is required to form the midbody after ring closure. We hypothesized that persistent Ect2^3A^ at the midbody could sustain RhoA activity, leading to an increase in ring components that prevent a proper midbody from being formed. To test this, we imaged endogenously tagged mNeonGreen-RhoA in HeLa cells rescued with Ect2 or Ect2^3A^ (Figure S4A). We saw RhoA remain at the midbody in cells rescued with Ect2^3A^ (Figure S4A). Next, we measured the changes in RhoA fluorescence at the midbody over time, using Ect2 as a marker (Figure S4B). While RhoA decreased at similar rates in cells rescued with Ect2 or Ect2^3A^, RhoA persisted at the midbody in Ect2^3A^ cells, although continued to decrease over time (Figure S4B). However, since active RhoA is a portion of the total RhoA pool, we visualized active RhoA more specifically (Figure 5A). To do this, we imaged cells expressing the RhoA biosensor (mNG x 2 RhoA-GTP binding domain from Rhotekin; rGBD) rescued with Ect2 or Ect2^3A^ (Figures 5B, C). As shown in Figure 5D, active RhoA appeared similar in Ect2 and Ect2^3A^ cells during ring assembly and ingression but then appeared broader and brighter at the midbody during the ring to midbody transition in Ect2^3A^ cells compared to Ect2 cells. We also observed active RhoA on either side of the unstable intercellular bridges similar to anillin (Figure 5E). We then quantified the levels of active RhoA in the midbody starting from ring closure and found that active RhoA persisted for a longer period of time in cells rescued with Ect2^3A^ compared to Ect2 (Figure 5F). Therefore, the persistence of Ect2 at the midbody caused by defective nuclear transport causes continuous RhoA activation at the midbody, which could cause defects in midbody formation and instability of the intercellular bridge.

**Figure 5.**
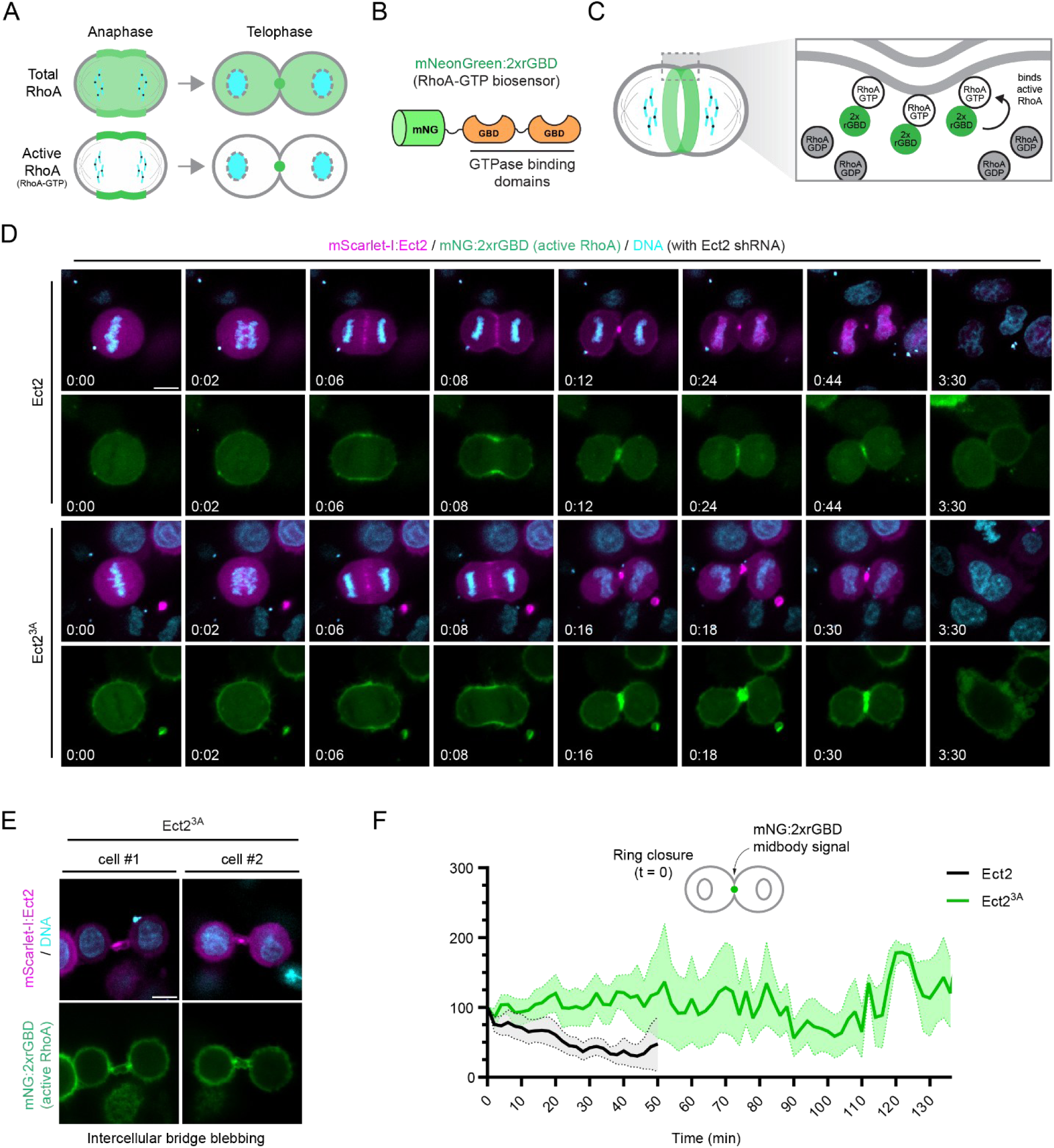
Ect2 nuclear localization is required to reduce active RhoA at the midbody and form a stable intercellular bridge. (A) Cartoon pictures show the localization of total RhoA (light green) in cells compared to active RhoA (dark green). Chromosomes are in blue. (B) A schematic shows the structure of the RhoA-GTP biosensor with mNeonGreen (mNG, green) fused to two RhoA-GTP binding domains (GBD, orange) from Rhotekin (Mahlandt et al., 2021). (C) A cartoon cell shows the localization of the biosensor (2xrGBD, green) for RhoA-GTP (white) at the equatorial cortex. RhoA-GDP is in grey. (D) Timelapse images show HeLa cells expressing RNAi-resistant mScarlet-I:Ect2 (magenta; Ect2 or Ect2^3A^), depleted of endogenous Ect2 using shRNA, co-expressing mNG:2xrGBD (green), and co-stained for DNA (cyan, Hoechst). The scale bar is 10 μm. Times are indicated in hours:minutes (t = 0 anaphase onset). (E) Images show intercellular bridge blebbing two Ect2^3A^ cells from D. The scale bar is 10 μm. (F) A line graph shows the normalized fluorescence intensity of mNG:2xrGBD (y-axis, a.u.) starting after ring closure (t = 0) for cells imaged as shown in D rescued with Ect2 (black, n = 9) or Ect2^3A^ (green, n = 8). The filled-in region surrounding the solid lines indicate standard error of the mean (SEM).

Ect2’s GEF activity is required to activate RhoA for cytokinesis, and we determined if the persistence of active RhoA is caused by Ect2’s GEF activity by mutating the DH domain, which is required for GDP to GTP exchange (Figures 6A, B). We predicted that if Ect2’s activation of RhoA caused midbody defects, then mutating the DH domain would prevent the dominant-negative effect caused by over-expression of the Ect2^3A^ mutant. Imaging cells with overexpressed Ect2^3A-LOF^ revealed that it decreased at the midbody over time similar to Ect2 (Figure 6C). Indeed, while 77.3% of cells with over-expressed Ect2^3A^ failed cytokinesis, only 8.3% cells failed cytokinesis with over-expressed Ect2^3A-LOF^, which was similar to Ect2 (16.6%; Figure 6D). As summarized in Figure 6E, these results suggest that when Ect2^3A^ persists at the midbody, it continues to activate RhoA, which prevents a functional midbody from forming.

**Figure 6.**
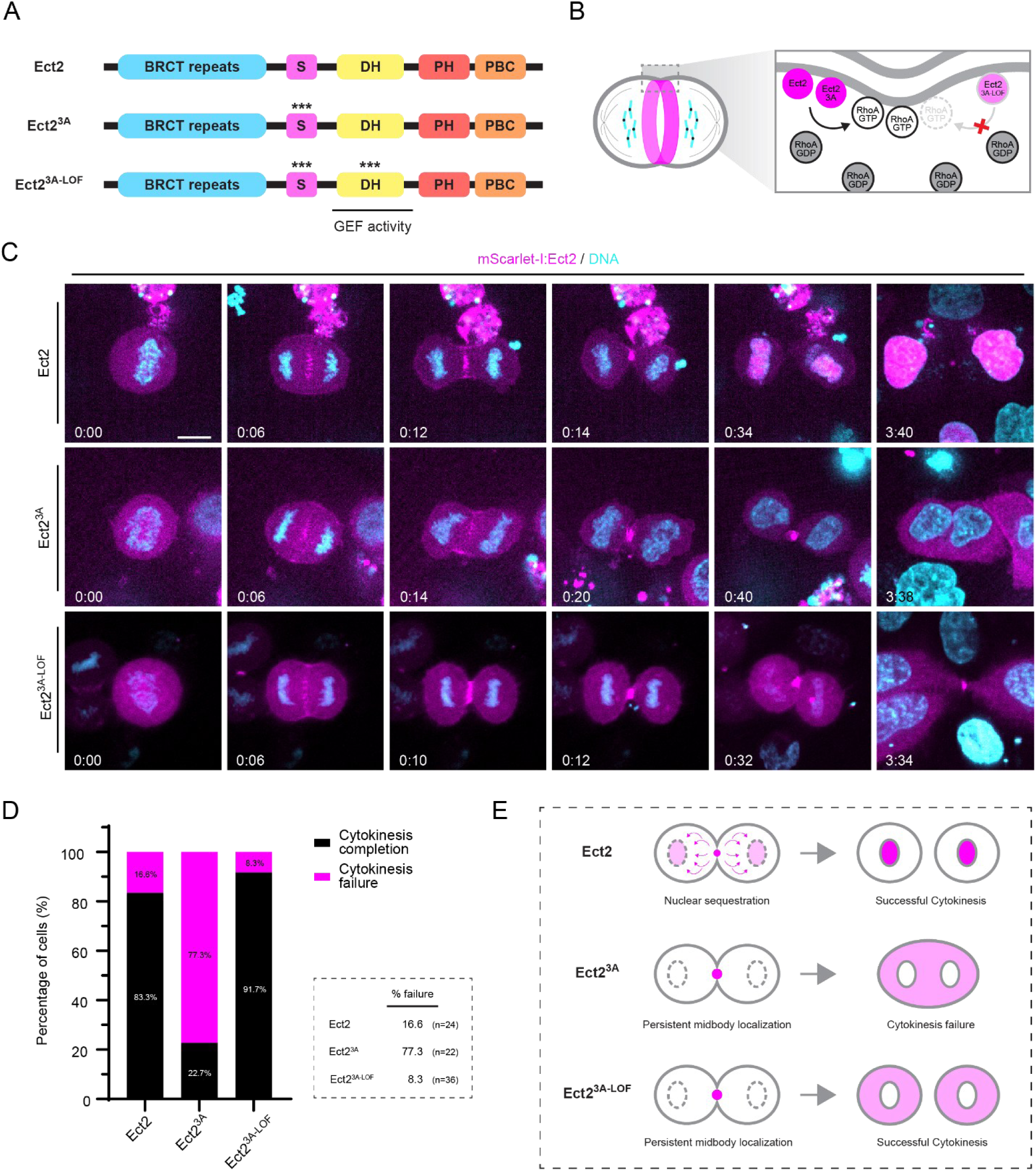
A decrease in Ect2 activity is required to form a stable midbody. (A) The structures of Ect2 with the location of the mutations (***) in the NLS and DH domain used in C-D are indicated for Ect2^3A^ and Ect2^3A-LOF^. (B) A cartoon cell shows how Ect2 and Ect2^3A^ (dark pink) have functional DH domains and can activate RhoA (RhoA-GTP, white), whereas Ect2^3A-LOF^ (light pink) cannot due to loss-of-function mutations in the DH domain. RhoA-GDP is shown in grey. (C) Timelapse images show HeLa cells expressing mScarlet-I:Ect2, Ect2^3A^ or Ect2^3A-LOF^ (magenta), co-stained for DNA (cyan, SYTO Deep Red). The scale bar is 10 μm. Time is in hours:minutes. (t = 0 is anaphase onset). (D) A bar graph shows the proportion of cells that succeed (black) or fail (pink) cytokinesis for cells treated as shown in C (Ect2 16.6% failure, n = 24; Ect2^3A^ 77.3% failure, n = 22; and Ect2^3A-LOF^ 8.3% failure, n = 36). (E) Cartoon cells show the changes in Ect2 localization (magenta) at the midbody and daughter cell nuclei when the NLS is mutated (Ect2^3A^) and both the NLS and DH domains are mutated Ect2^3A-LOF^.

## Discussion

Our study shows that the nuclear transport of Ect2 is required for its removal from the midbody for cytokinesis. For the first time, we use an endogenous tag to show the dynamic changes in Ect2 localization throughout cytokinesis, until abscission. We also mutated the NLS in the S-loop region and revealed that this site is required for cytokinesis in rescue assays, and causes dominant-negative effects when over-expressed in both HeLa and HEK293T cells. When the NLS is mutated, Ect2 remains at the midbody in cells, causing intercellular bridge instability and membrane regression. Using a RhoA biosensor, we showed that Ect2 continuously activates RhoA at the midbody and intercellular bridge, and cytokinesis can be restored by localizing the NLS mutant to the nucleus, or by inactivation of the DH domain, which is required for nucleotide exchange to generate active RhoA. Thus, we propose that Ect2 must be removed from the midbody to prevent the continuous activation of RhoA for the transition to machinery required for midbody maturation and abscission (Figure 7).

**Figure 7.**
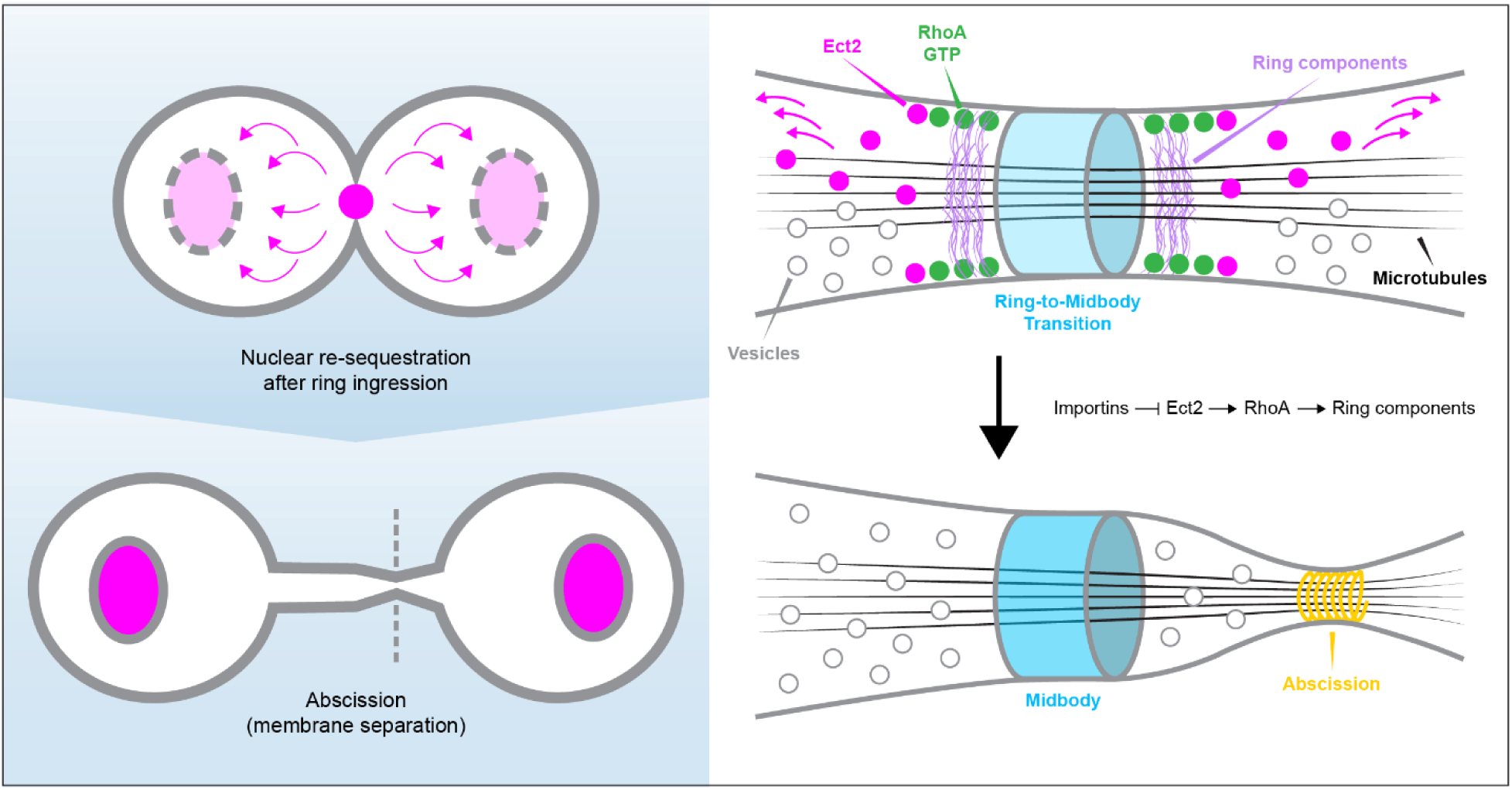
A model for nuclear sequestration as a regulatory mechanism for Ect2 function in cytokinesis. After ring ingression and closure, Ect2 (pink) is removed from the midbody (blue) via nuclear transport by importins to reduce RhoA-GTP (green) at the intercellular bridge. A reduction in active RhoA is likely important to decrease ring components (purple) for intercellular bridge stability and midbody maturation for abscission (yellow). When the NLS of Ect2 is mutated it remains at the midbody and active RhoA persists causing intercellular bridge blebbing and cytokinesis failure, likely by sustaining ring components such as actin and myosin. This mechanism spatiotemporally couples ring closure with nuclear envelope reformation, and could regulate other contractile proteins.

Previous work suggested that the levels of Ect2 the midbody is important for abscission. Ect2 reduction at the midbody was proposed to be required to permit Fip3 to form a complex with Cyk4. This was supported by binding studies where Ect2 competes with Fip3 for Cyk4-binding. (Wilson et al., 2005; Simon et al., 2008). However, due to the lack of tools for live-cell imaging of Ect2, these studies relied on antibodies making it difficult to precisely stage cells. Similarly, overexpression of the N-terminus of Ect2 (which binds to Cyk4) causes cytokinesis failure, although the reason for this failure was not clear (Chalamalasetty et al., 2006). Our studies show that the timing of Ect2’s removal from the midbody happens much earlier than Fip3’s localization at the midbody. Therefore, while the persistence of Ect2 at the midbody likely eventually leads to defects that include blocking Fip3 localization, the primary defect occurs earlier during the ring to midbody transition.

Although the midbody is an extremely dense structure, the removal of Ect2 occurs soon after ingression, likely before a stable midbody has fully formed. However, midbody assembly is not well-understood, and it is not clear when it becomes a more stable, dense structure. Presumably, Ect2 retains interactions with phospholipids in the overlying membrane, which could prevent Ect2 from being integrated into the denser regions of the midbody.

We hypothesize that nuclear re-sequestration acts as a transition mechanism between ring closure and midbody formation, by reducing RhoA-dependent ring proteins. This elegant mechanism begins to remove Ect2 and reduce RhoA activity as early as nuclear envelope reformation, which coincides with ring closure. This regulation may be faster than relying on mechanisms such as degradation or inhibition by post-translational modifications, which requires the synthesis and/or spatiotemporal control of specific enzymes. Since we showed that Ect^3A^ causes cytokinesis failure in both HeLa and HEK293T cells, this mechanism is potentially conserved.

Other cytokinesis proteins, such as anillin, Cyk4, Mklp1 and mDia2 have NLSs, and it is exciting to speculate that a similar mechanism could control other cytokinesis proteins either during or post cytokinesis. Overexpression of an NLS mutant of anillin in HeLa cells does not lead to cytokinesis failure, but causes cell shape changes in interphase, indicating that the nuclear sequestration of anillin maintains integrity of the cytoskeleton (Chen et al., 2015). Interestingly, overexpression of an NLS mutant of Mklp1/Kif23 causes cells to remain connected by an intercellular bridge suggesting a block in abscission (Liu and Erikson, 2007). How Mklp1 (or excess Mklp1) at the midbody causes abscission failure is unclear. Unlike Ect2, Mklp1 remains at the midbody for a long time, and the timing of its localization to the nuclei has not been studied in live cells (Makyio et al., 2012). Phosphorylation of the NLS in Mklp1 attenuates nuclear localization, and a phosphodeficient mutant leads to premature nuclear localization after ring closure and causes cytokinesis failure, indicating that nuclear transport is controlled by phosphorylation (Neef et al., 2006; Liu and Erikson, 2007).

We also found a minor delay in the duration of ingression and impairment of ring closure in cells rescued with the Ect2 NLS mutant. Slower ingression kinetics could be due to the Ect2 mutant being less efficient at generating contractility through RhoA. However, this was not supported by our data showing that the localization of anillin and active RhoA was similar to the control. However, Ect2 itself was more variable, and it is possible that importin-binding to the S-loop affects other functions of Ect2 that could impact ingression independent of active RhoA. Furthermore, there could be small changes in active RhoA that are not measurable by the biosensor.

Ect2 is a proto-oncogene and is overexpressed in various cancers (Justilien et al., 2019). The overexpression of a C-terminal fragment containing a mutant NLS confers transformation to HEK293 cells (Saito et al., 2004). Other studies showed that mis-localization of Ect2 in the cytosol is associated with a poor prognosis in cancer patients with lung and colorectal cancers (Kosibaty et al., 2019; Cook et al., 2021; Yi et al., 2022). The mechanism for Ect2 in promoting tumorigenesis is believed to be through its function in activating both cytosolic and nuclear Rac1, resulting in an increase in pathways that include ribosomal biogenesis, cell proliferation and metastasis (Fields and Justilien, 2010). Our data suggests that mis-localized Ect2 in cancer cells has the potential to cause cytokinesis failure, which can increase chromosomal instability, an important cancer hallmark. Whole genome-duplication by cytokinesis failure can contribute to aneuploidy, when binucleated cells avoid cell death and divide with lagging chromosomes, multiple poles or cleavage furrows (Lens and Medema, 2019). This work provides a new mechanism for how Ect2 could promote tumorigenesis.

---

**Supplementary Figure 1.**
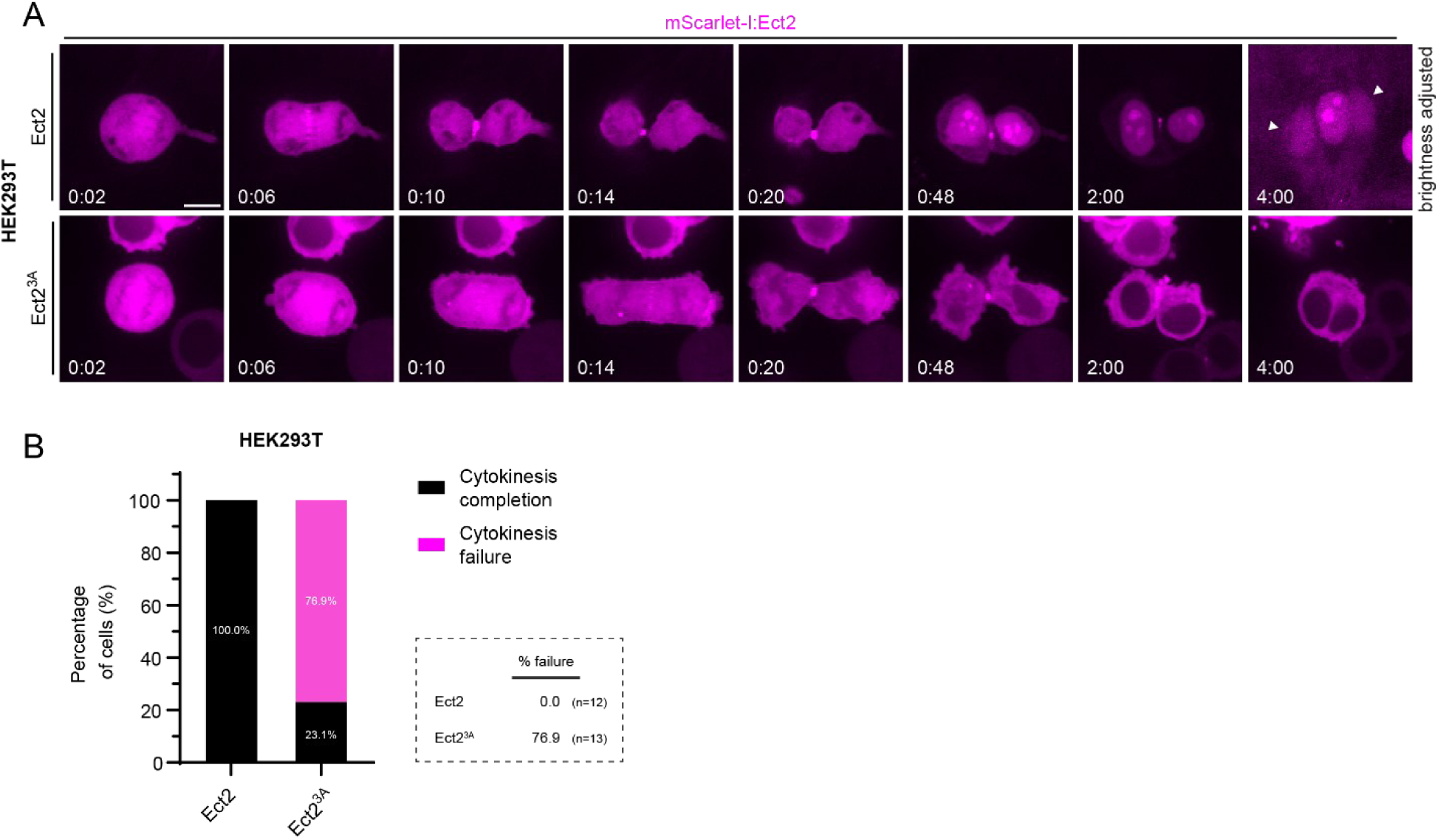
The NLS of Ect2 is required for cytokinesis in HEK293T cells. (A) Timelapse images show HEK293T cells expressing mScarlet-I-Ect2 or Ect2^3A^ (magenta). The scale bar is 10 μm. Time is in hours:minutes (t = 0 is anaphase onset). The contrast was adjusted for the last panel (t = 4:00) of Ect2 due to low signal. White arrows point to the two nuclei of the original dividing cell. (B) A bar graph shows proportion of cells that succeed (black) or fail (pink) cytokinesis for cells treated as in A (Ect2 0% failure, n = 12; Ect2^3A^ 76.9% failure, n = 13).

**Supplementary Figure 2.**
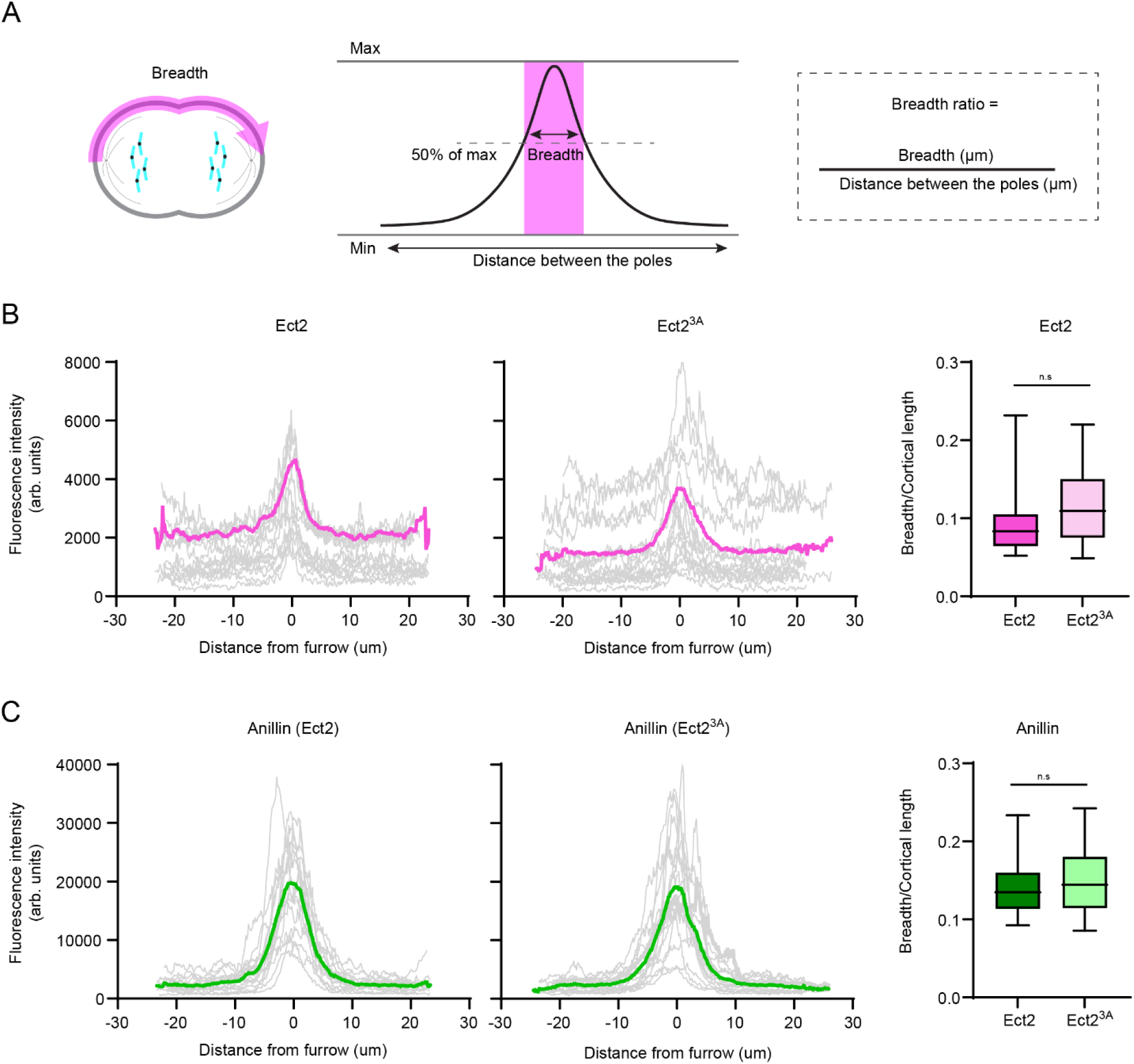
The NLS of Ect2 is not required for the localization of Ect2 or anillin at the equatorial cortex. (A) A schematic shows how breadth of accumulated Ect2 or anillin the equatorial cortex was measured using line scans. A line was drawn along half of the perimeter of a cell in early ingression to measure the intensity of pixels along the line. The breadth was defined as the number of pixels >50% of the normalized maximum intensity, which was converted into μm. To control for cell size, the breadth was divided by the length. (B-C) Graphs (left) show fluorescence intensity (y-axis, a.u.) over distance from the ingressing cortex (x-axis, point of ingression is 0, μm) for (B) mScarlet-I:Ect2 or Ect2^3A^ co-depleted for Ect2 with shRNAs in HeLa cells with (C) endogenous mNeonGreen:anillin (n = 17 for Ect2 and Ect^3A^). To the right, box-and-whiskers plots show the ratio of the breadth divided by the cortical length. Statistical analysis was performed using a Student’s T-test for mNeonGreen:anillin and non-parametic test for mScarlet-I:Ect2, and no statistically significant differences between the means were detected.

**Supplementary Figure 3.**
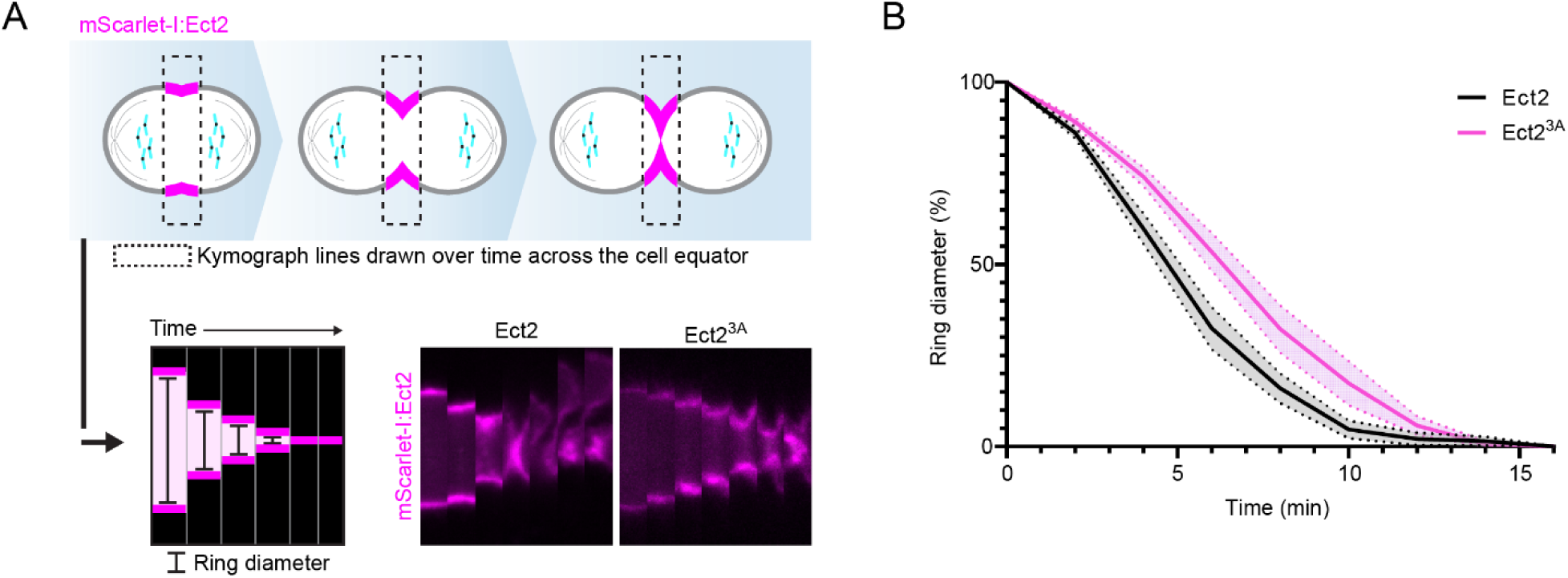
The NLS of Ect2 plays a minor role in ring ingression. (A) Cartoon cells shows the region that was used to generate kymographs for cells expressing mScarlet-I:Ect2 or Ect2^3A^ (magenta) as shown. (B) The line graph shows % change in ring diameter over time in cells expressing mScarlet-I:Ect2 (black, n = 24) or Ect2^3A^ (pink, n = 13) depleted for endogenous Ect2 with shRNA. The bars show standard error of the mean (SEM).

**Supplementary Figure 4.**
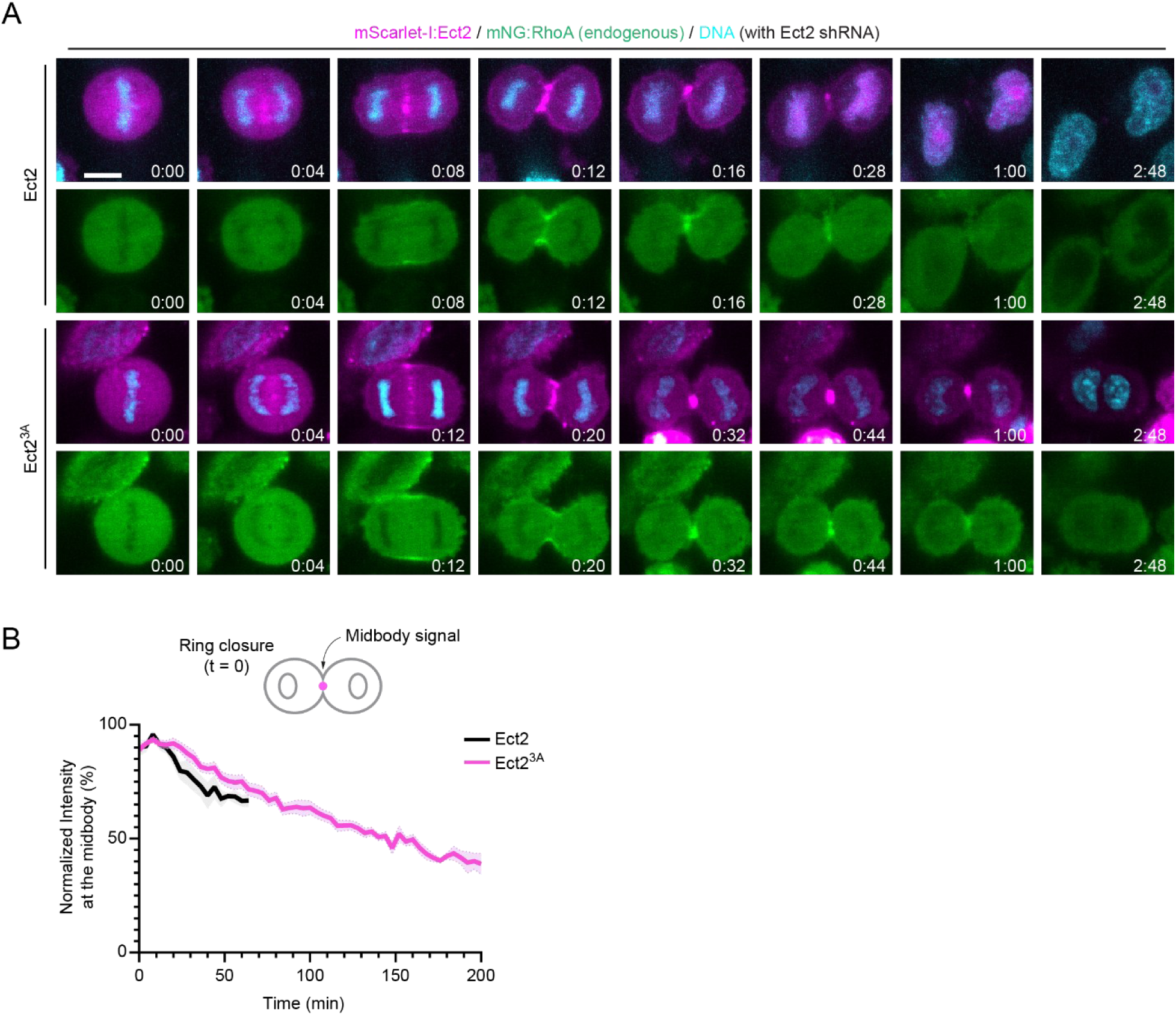
Total RhoA at the midbody decreases at a rate that is similar for Ect2 and NLS mutant Ect2. (A) Timelapse images show HeLa cells with endogenous mNeonGreen:RhoA (green), co-expressing mScarlet-I:Ect2 or Ect2^3A^ (magenta), depleted for Ect2 with shRNA and co-stained for DNA (cyan, Hoechst). The scale bar is 10 μm. Time is in hours:minutes. (t = 0 is anaphase onset). (B) A line graph shows the normalized total fluorescence intensity (y-axis, %) of mNeonGreen:RhoA at the midbody in Ect2 (n = 10) or Ect2^3A^ cells (n = 15) over time (x-axis, minutes; ring closure t = 0). The error bars show standard error of the mean (SEM).

